# Clones and Clusters of Antimicrobial-Resistant *Klebsiella* from Southwestern Nigeria

**DOI:** 10.1101/2021.06.21.449255

**Authors:** Ayorinde O. Afolayan, Anderson O. Oaikhena, Aaron O. Aboderin, Olatunde F. Olabisi, Adewale A. Amupitan, Oyekola V. Abiri, Veronica O. Ogunleye, Anthony Underwood, Erkison Ewomazino Odih, Abolaji T. Adeyemo, Adeyemi T. Adeyemo, Temitope O. Obadare, Sophia David, Silvia Argimón, Monica Abrudan, Abiodun Egwuenu, Chikwe Ihekweazu, David M. Aanensen, Iruka N. Okeke, the NIHR Global Health Research Unit (GHRU) on Genomic Surveillance of Antimicrobial Resistance

## Abstract

**Introduction:** *Klebsiella pneumoniae* is a World Health Organization high-priority antibiotic-resistant pathogen. However, little is known about the population structure and evolution of *Klebsiella* circulating in Nigeria.

**Methods:** We performed whole genome sequencing (WGS) of 141 *Klebsiella* isolated between 2016 and 2018 from clinical specimens at 3 antimicrobial-resistance (AMR) sentinel surveillance tertiary hospitals in southwestern Nigeria. We conducted *in silico* multilocus sequence typing, AMR gene, virulence gene, plasmid, and K and O loci profiling, as well as phylogenetic analyses, using publicly available tools and Nextflow pipelines.

**Results:** Phylogenetic analysis revealed that the majority of the 134 *K. pneumoniae* and 5 *K. quasipneumoniae* isolates from Nigeria characterized are closely related to globally disseminated multidrug-resistant clones. Of the 39 *K. pneumoniae* sequence types (STs) identified, the most common were ST307 (15%), ST5241 (12%), ST15 (~9%), and ST25 (~6%). ST5241, one of 10 novel STs detected, is a single locus variant of ST636 carrying *dfrA14*, *tetD*, *qnrS*, and *oqxAB* resistance genes. The extended-spectrum β lactamase (ESBL) gene *bla*CTX_M-15 was seen in 72 % of *K. pneumoniae* genomes, while 8% encoded a carbapenemase. Four likely outbreak clusters from one facility, within STs 17, 25, 307, and 5241, were ESBL but not carbapenemase-bearing clones.

**Conclusion:** This study uncovered known and novel *K. pneumoniae* lineages circulating in Nigeria that include multidrug-resistant ESBL producers. Carbapenemase-producing isolates remain uncommon. WGS retrospectively identified outbreak clusters, pointing to the value of genomic approaches in AMR surveillance for improving infection prevention and control in Nigerian hospitals.

**summary:** We performed whole genome sequencing (WGS) of 141 *Klebsiella* isolated in 2016-2018 at 3 antimicrobial-resistance (AMR) sentinel surveillance tertiary hospitals in southwestern Nigeria. This study uncovered known and novel *K. pneumoniae* lineages circulating in Nigeria that include multidrug-resistant ESBL producers.

**FUNDING:** This work was supported by Official Development Assistance (ODA) funding from the National Institute of Health Research [16/136/111: NIHR Global Health Research Unit on Genomic Surveillance of Antimicrobial Resistance].

This research was commissioned by the National Institute of Health Research using Official Development Assistance (ODA) funding. INO is an African Research Leader supported by the UK Medical Research Council (MRC) and the UK Department for International Development (DFID) under the MRC/DFID Concordat agreement that is also part of the EDCTP2 program supported by the European Union. The funders had no role in the content, crafting or submission of this paper. The views expressed in this publication are those of the authors and not necessarily those of the funders or their affiliates.

**CONFLICT OF INTEREST:** The authors: No reported conflicts of interest. All authors have submitted the ICMJE Form for Disclosure of Potential Conflicts of Interest.

## INTRODUCTION

*Klebsiella* is a ubiquitous gram-negative genus that can cause a variety of opportunistic infections [1, 2]. *Klebsiella pneumoniae,* the species most commonly associated with human disease, is frequently implicated in life-threatening bacteremia and sepsis arising from translocations from non-sterile niches or medical devices in hospitals [3]. Although the precise burden of infectious diseases resulting from *Klebsiella* spp. is unknown in Africa, there is an upward trend in reports of *K. pneumoniae*-associated bloodstream infections, including from Nigeria, Côte d’Ivoire, Malawi, Gambia, and South Africa [4–8]. There have also been isolated case reports of non-*pneumoniae* infections in Nigeria [9]. Altogether, these infections have high mortality rates and incur high costs [10, 11].

Rigorous surveillance of *Klebsiella* clones circulating within Nigerian hospitals and communities is needed to inform treatment guidelines and prioritize AMR interventions. Whole genome sequencing (WGS) offers the attractive prospect of meeting this need within local resource constraints. WGS tools also provide taxonomic resolution below serotype levels, a feat impossible in resource-limited settings by classical typing methods [13]. Mapping the diversity of *Klebsiella* populations at high resolution, with spatiotemporal dynamics, also makes it possible to elucidate the origin and dissemination of AMR [12, 13].

In contrast to similarly constrained settings elsewhere in Africa [14–16], few Nigeria reports include in-depth analysis of more than a handful of *Klebsiella* isolates. For example, as of the time of writing, Pathogenwatch, which sources its public genomes from ENA, includes only 109 *Klebsiella* genomes from Nigeria [17]. Nigeria recently initiated national AMR surveillance following commissioning of a 2017 National Action Plan, and the Nigeria Centre for Disease Control (NCDC) enrolled the country in the WHO’s Global Antimicrobial Resistance Surveillance System (GLASS) [18]. NCDC is building a network that currently consists of 9 AMR sentinel hospital laboratories and 2 national reference laboratories [19]. The very low number of *Klebsiella* genomes from Nigeria compelled the Nigeria node of the Global Health Research Unit (GHRU) on Genomic Surveillance of AMR, which provides WGS services to the reference laboratories, to collect and sequence retrospective isolates. In addition to increasing the number of relevant genomes in publicly available databases that can support prospective surveillance, this study aimed to understand the population structure, evolution, pathogenicity, and transmission dynamics of this critical-priority pathogen circulating in southwestern Nigeria.

## METHODS

### Ethical Considerations

Most genomes included in this study were acquired from surveillance without linked patient data. We additionally integrated data from the Obafemi Awolowo University (OAU) Teaching Hospital sentinel site, as approved by the Ethics and Research Committee, Obafemi Awolowo University Teaching Hospitals Complex with registration number ERC/2017/05/06.

### GHRU Nigeria Workflow

#### Strains, re-identification, and antimicrobial susceptibility testing

The NCDC, Nigeria’s AMR surveillance coordinating center, requested from the network cryopreserved invasive isolates (from blood, cerebrospinal fluid, and urine) from 2016-2018, to develop a local genome database. Sentinel laboratories provided the national reference laboratory with strains and information on species identity, clinical diagnosis, and antimicrobial susceptibility tests (AST) via WHONET [20]. At the national reference laboratory, in collaboration with the GHRU, isolate identification and AST were validated using the VITEK system (version 2.0), testing, as appropriate, amikacin, gentamicin, ampicillin, amoxicillin/clavulanic acid, piperacillin/tazobactam, cefuroxime, cefuroxime_axetil, cefepime, ceftriaxone, cefoperazone/sulbactam, nitrofurantoin, nalidixic acid, ciprofloxacin, ertapenem, imipenem, meropenem, trimethoprim-sulphamethoxazole, colistin, and tigecycline [21]. Antimicrobial susceptibility test results were interpreted in line with the CLSI standards [22].

#### DNA extraction, Library Preparation, and Sequencing

Genomic DNA was extracted using the Wizard DNA extraction kit (Promega; Wisconsin, USA; Cat. No: A1125). DNA was quantified using a dsDNA Broad Range fluorometric quantification assay (Invitrogen; California, USA; Cat. No: Q32853). Double-stranded DNA libraries (avg. 500 bp) were prepared using the Covaris LC220 for fragmentation, and NEBNext Ultra II FS DNA library kit for Illumina with 384-unique indexes (New England Biolabs, Massachusetts, USA; Cat. No: E6617L). Libraries were sequenced using the HiSeq X10 with 150 bp paired-end chemistry (Illumina, CA, USA).

#### Genome assembly

Genome assembly was carried out according to the GHRU protocol (https://www.protocols.io/view/ghru-genomic-surveillance-of-antimicrobial-resista-bpn6mmhe). Parameters for post-assembly quality checks include the total genome size (between 4584497 bp to 7012008 bp), N50 score (> 25000), contaminant level (< 5%), and number of contigs (< 300). Only *Klebsiella* genomes that passed all quality checks (*K. pneumoniae*, n = 134; *K. quasipneumoniae*, n = 5) were selected for downstream analysis.

### Single Nucleotide Polymorphism (SNP) Analysis

Sequence reads were mapped to the reference genome of the *K. pneumoniae* strain NTUH-K2044 (Genbank Accession number <u>GCF_009497695.1</u>) to determine evolutionary relationships among the isolates according to GHRU protocol (https://www.protocols.io/view/ghru-genomic-surveillance-of-antimicrobial-resista-bpn6mmhe). Pairwise SNP distances for likely outbreak isolates from OAU were calculated from the pseudo-genome alignment using FastaDist (https://gitlab.com/antunderwood/fastadist). Outbreak isolates belonging to STs 17, 25, and 307 were aligned to reference genomes NZ_CP009461.1, NZ_CP031810.1, and NZ_CP022924.1 respectively, selected with Bactinspector (https://gitlab.com/antunderwood/bactinspector/). The ST636 reference EuSCAPE_UK136 was selected from Pathogenwatch collections, being the only known single locus variant of ST5241.

### Prediction of AMR Determinants, Virulence Factors, and Plasmid Profiling

AMR determinants and plasmid replicons were predicted in accordance with the aforementioned GHRU protocol. *omp* mutations, virulence score, and acquired virulence genes were determined using Kleborate (v2.0.0; https://github.com/katholt/Kleborate) [23, 24].

### Concordance Analysis

Concordance between phenotypic and genotypic AMR for aminoglycosides, β-lactams, carbapenem, cephalosporin, quinolone, and trimethoprim was determined by using https://gitlab.com/-/snippets/2050300 based on a protocol (https://glcdn.githack.com/cgps/ghru/ghrur/raw/master/docs/articles/simple_concordance_example.html) that employs the epi.tests function of the epiR package to calculate sensitivity and specificity with confidence bounds.

### Identification of Multilocus Sequence Types

Multilocus sequence types (MLSTs) according to the Pasteur scheme were determined by following the aforementioned GHRU protocol (https://www.protocols.io/view/ghru-genomic-surveillance-of-antimicrobial-resista-bpn6mmhe) [25]. We also used Kleborate (v2.0.0; https://github.com/katholt/Kleborate) to identify the ST most closely related to novel STs (in terms of locus variants).

### Availability of Sequence Data

Raw sequence datasets generated during this study have been deposited in the ENA with bioproject number PRJEB29739 (https://www.ebi.ac.uk/ena/browser/view/PRJEB29739). Accessions are available in Supplementary Table 3.

## RESULTS

### Epidemiology, Sequence Types, Virulence-Associated Determinants, and AMR Determinants of *Klebsiella* from Southwestern Nigeria Hospitals

Retrospective *Klebsiella* isolates were collected from 3 hospitals that had archival isolates: Obafemi Awolowo University Teaching Hospital (OAU, Ile-Ife, Osun State, Nigeria; n = 92), University College Hospital (UCH, Ibadan, Oyo State, Nigeria; n = 30), and Ladoke Akintola University Teaching Hospital (LAU, Oshogbo, Osun State, Nigeria; n = 12). Of the 134 isolates, 87 were from blood, comprising 45 from OAU, 30 from UCH, and 12 from LAU. We additionally sequenced 47 non-blood isolates from OAU, recovered from urine (n = 37), ocular (n = 2), stool (1), throat (n = 2), and rectal (n = 5) swabs.

Of the 134 genomes confirmed as *K. pneumoniae* by WGS, 18 (13%) and 33 (25 %) were identified by reference laboratory VITEK and the sentinel laboratories, respectively, as either other members of the *Enterobacteriaceae* family, *Acinetobacter baumannii*, or *Pseudomonas aeruginosa* (Supplementary Table 1). All WGS-identified *Klebsiella quasipneumoniae* isolates were identified either as *K. pneumoniae* or *Enterobacter* complexes by VITEK and the sentinel laboratories, respectively (Supplementary Table 1). The phylogenetic tree, epidemiological data, and analyses data can be visualized in Microreact for retrospective *Klebsiella pneumoniae* (https://microreact.org/project/GHRUNigeriaKpneumoniae/2a856694), and OAU outbreak isolates (ST17: https://microreact.org/project/brfq17BXzwmqptfcrNRfZR/71b2f0f1; ST25: https://microreact.org/project/8FP4F1D5fSQMv6FDxbR39b/bd381f0b; ST307: https://microreact.org/project/sV5NsJ8szcorvFAeFQgV4E/6498edf1; ST5241: https://microreact.org/project/u5RPX7CjyitjRYMQ9ByW4W/1a3570e2).

Overall, the *K. pneumoniae* isolates sequenced in this study belonged to 39 different sequence types (STs), with ST307, ST5241, ST15, and ST25 being most common (Figure 1). Only 4 STs (ST15, ST101, ST147, and ST307) were found across the 3 sentinel sites (Figure 1). *Klebsiella quasipneumoniae* genomes (n = 5) belonged to 4 STs: ST1602, ST2133, ST5250 (n = 1 each), and ST5249 (n = 2).

**Figure 1.**
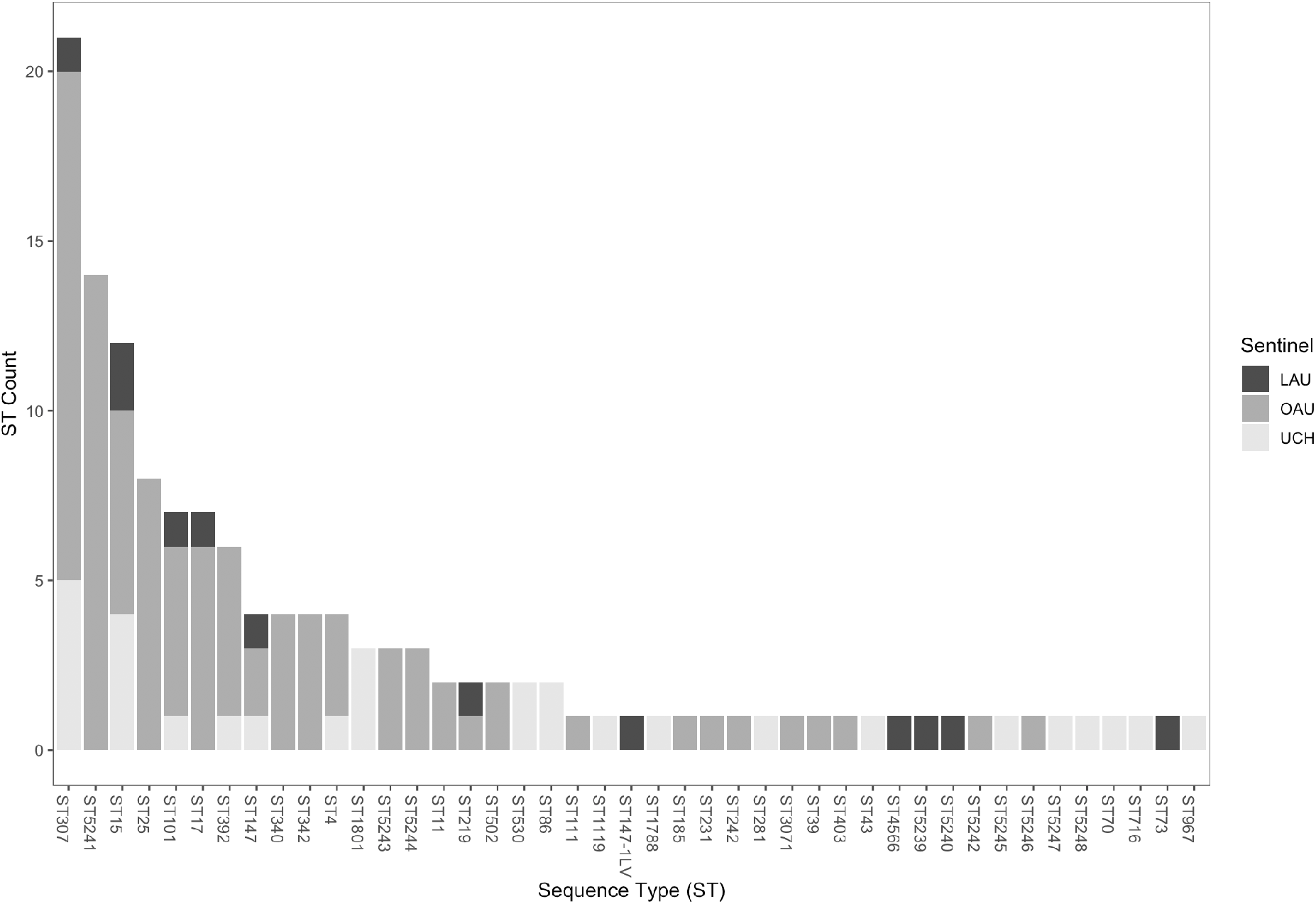
Sequence Types of *Klebsiella pneumoniae* and their distribution across the 3 sentinel sites. Key (LAU = Ladoke Akintola University Teaching Hospital, OAU = Obafemi Awolowo University Teaching Hospital, UCH = University College Hospital).

We identified few virulence genes associated with invasiveness, including yersiniabactin (*ybt* genes and *fyuA*; n = 42, 31.34%), aerobactin (*iuc*; n = 2), salmochelin (*iro*; n = 2), and capsule expression upregulators *rmpADC* and *rmpA2* (n = 2). Only 2 ST86 isolates had a virulence score of 3, the highest recorded in our collection. No acquired *K pneumoniae* virulence gene was detected in novel ST5241 (Figure 2), nor in the non-pneumoniae *Klebsiella* species.

**Figure 2.**
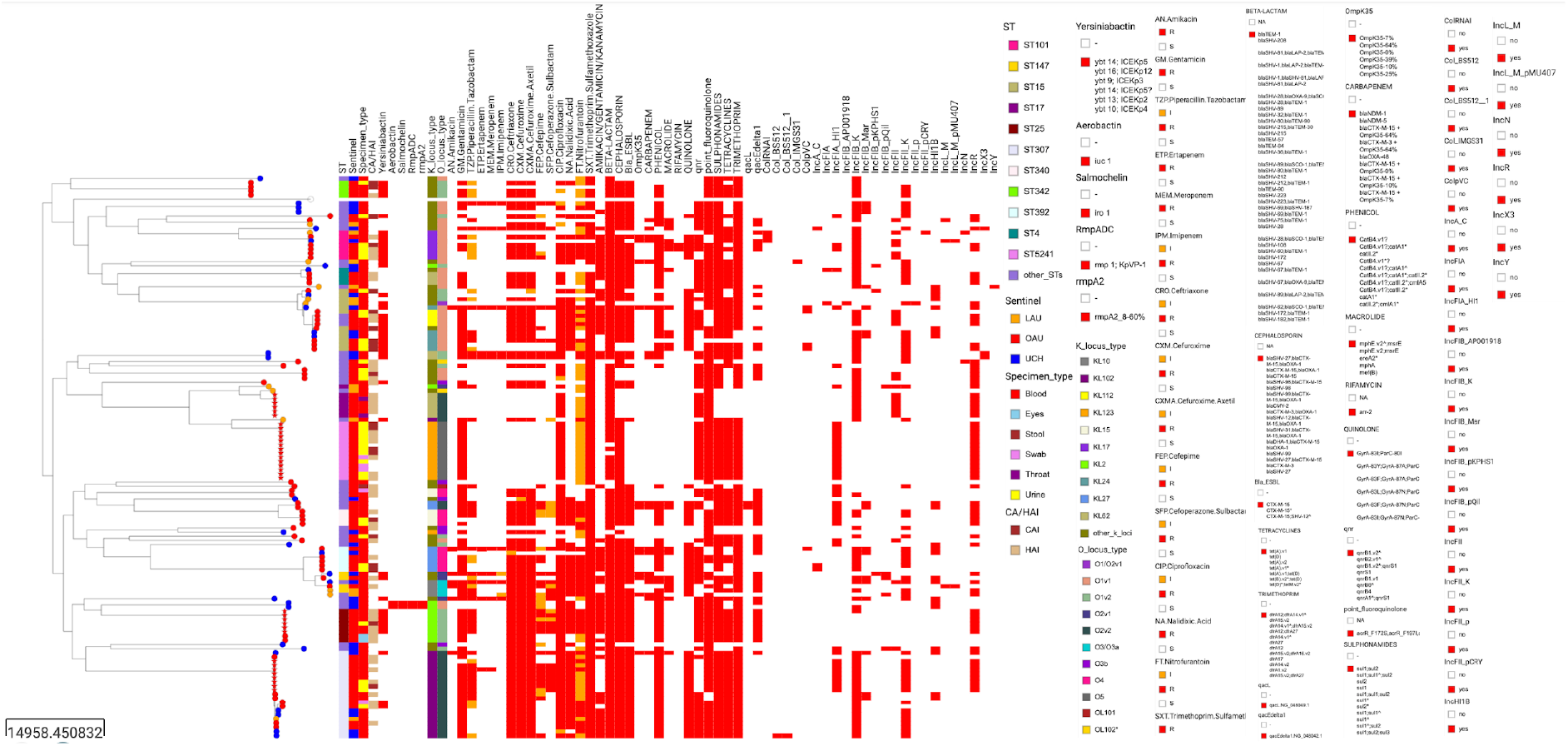
Epidemiological data, virulence determinants, antibiotic profile (phenotypic resistance), and antimicrobial resistance determinants in *K. pneumoniae* genomes ordered by phylogeny. The heat map shows the presence (red) or absence (blank) of virulence determinants, phenotypic resistance, AMR genes, and plasmid replicon genes. Tree nodes represent the origin of isolate collection. The shape of the tree nodes depicts the outbreak (star) and non-outbreak (circle) isolates. The data are available at https://microreact.org/project/GHRUNigeriaKpneumoniae/2a856694.

We detected 33 different phenotypically-defined K (capsular) loci from the invasive *K. pneumoniae* isolates sequenced, with KL102, KL123, KL2, KL62, and KL27 representing the 5 most common K loci (Supplementary Table 2). Eleven different O loci were detected among the invasive *K. pneumoniae* isolates recovered from bloodstream infections. Collectively, the 3 O loci – O1v1, O2v2, and O1v2 – accounted for more than 65% of the *K. pneumoniae* strains, and O5 was solely detected in ST5241 genomes.

Genes conferring resistance or reduced susceptibility to at least 5 antibiotic classes were detected in 116 (86.6%) of *K. pneumoniae* genomes. Aside from core *bla_SHV_*, these included8 β-lactamase genes, of which the extended-spectrum β-lactamase (ESBL) *bla_CTX-M-15_* was by far the most common, present in 71.6% (n = 96) of the *K. pneumoniae* isolates belonging to 33 STs (Figure 2). As seen in other studies, we found this ESBL in strains with plasmids belonging to IncFIB_K (n = 70/96), IncFII_K (n = 51/96), and IncR (n = 43/96) incompatibility groups [26, 27]. The *bla_NDM-1_* (n = 8/134), *bla_NDM-5_* (n = 2/134), and *bla_OXA-48_* (n = 1/134) genes were the only carbapenemases detected, and these occurred in 8 STs, including ST392 (*bla_NDM-1_*; n = 2), ST147/147-1-locus-variant (*bla_NDM-1_*; n = 2), ST530 (*bla_NDM-_ 5*; n = 2), ST43, ST15, ST307, ST716 (*bla_NDM-1_*; n = 1 each), and ST219 (*bla_OXA-48_*; n = 1) (Figure 2). OmpK35 truncations were observed together with *bla_CTX-M-15_* in 11 isolates and with *bla_NDM-1_* in 5 isolates. Altogether, 18 (13.5%) *K. pneumoniae* strains carried carbapenemase genes, and/or ESBL genes plus porin defects known to confer carbapenem resistance [28]. Phenotypic antimicrobial susceptibility testing found 11 of 128 isolates carbapenem non-susceptible, and *bla_NDM-1_* (n = 6), *bla_NDM-5_* (n = 2), and/or *bla_CTX-M-15_* and OmpK35 truncations (n = 1) could account for non-susceptibility in 9 of them (Figure 2). Concordance analysis showed that phenotypic AST and genotypic AMR gene prediction results agree substantially for carbapenems (concordance: 92.2%; sensitivity: 82% [48.22% - 97.72%]; specificity: 93.2% [86.97% - 97%]).

Reduced susceptibility to nalidixic acid (n = 70/125) or ciprofloxacin (n = 114/125) could be explained by the presence of one or more combinations of mutations in the quinolone resistance determining region (QRDR) of *gyrA* and *parC* (n = 60), plasmid-mediated quinolone resistance genes (*qnrS*, *qnrA*, *qnrB*; n = 82), and/or efflux-mediating mutations in *acrR* gene (n = 98) (Figure 2). The 5 *K. quasipneumoniae* were resistant to phenicols, tetracyclines, sulphonamides, trimethoprim, and β-lactams (https://microreact.org/project/4f6Y56ECE979wkj8oDBhk2/dee16dac). One LAU isolate, collected 6 months after an outbreak of *K. quasipneumoniae* strains carrying *bla_NDM-5_* in Abuja, Nigeria, was remotely related to those outbreak strains (SNP distance ≥ 235), but, like the other 4 *K. quasipneumoniae* from the current study, it lacked carbapenemase genes [9].

### Retrospective Characterization of Possible Outbreaks

Recovery dates for 74 of 92 isolates from the OAU facility were supplied. In under a year, between 6 and 14 isolates were recovered belonging to ST17, ST25, ST307, and ST5241. We noticed clusters of closely related isolates that were recovered within short periods, and we hypothesized that these might represent unreported/undetected outbreaks. Intra-cluster pairwise distances within the potential outbreak clades ranged from 1 to 3 SNPs, and inter-cluster pairwise distances within the same ST ranged from 46 to 370. Temporally-clustered, closely-related clonal groups within each of the four STs (Figure 3), with similar AMR gene, virulence gene, and plasmid replicon profiles, support that these clusters represent outbreaks, and that nosocomial transmission contributes heavily to invasive *Klebsiella* infections. The outbreaks mostly involved ESBL lineages but no carbapenemase-producers.

**Figure 3:**
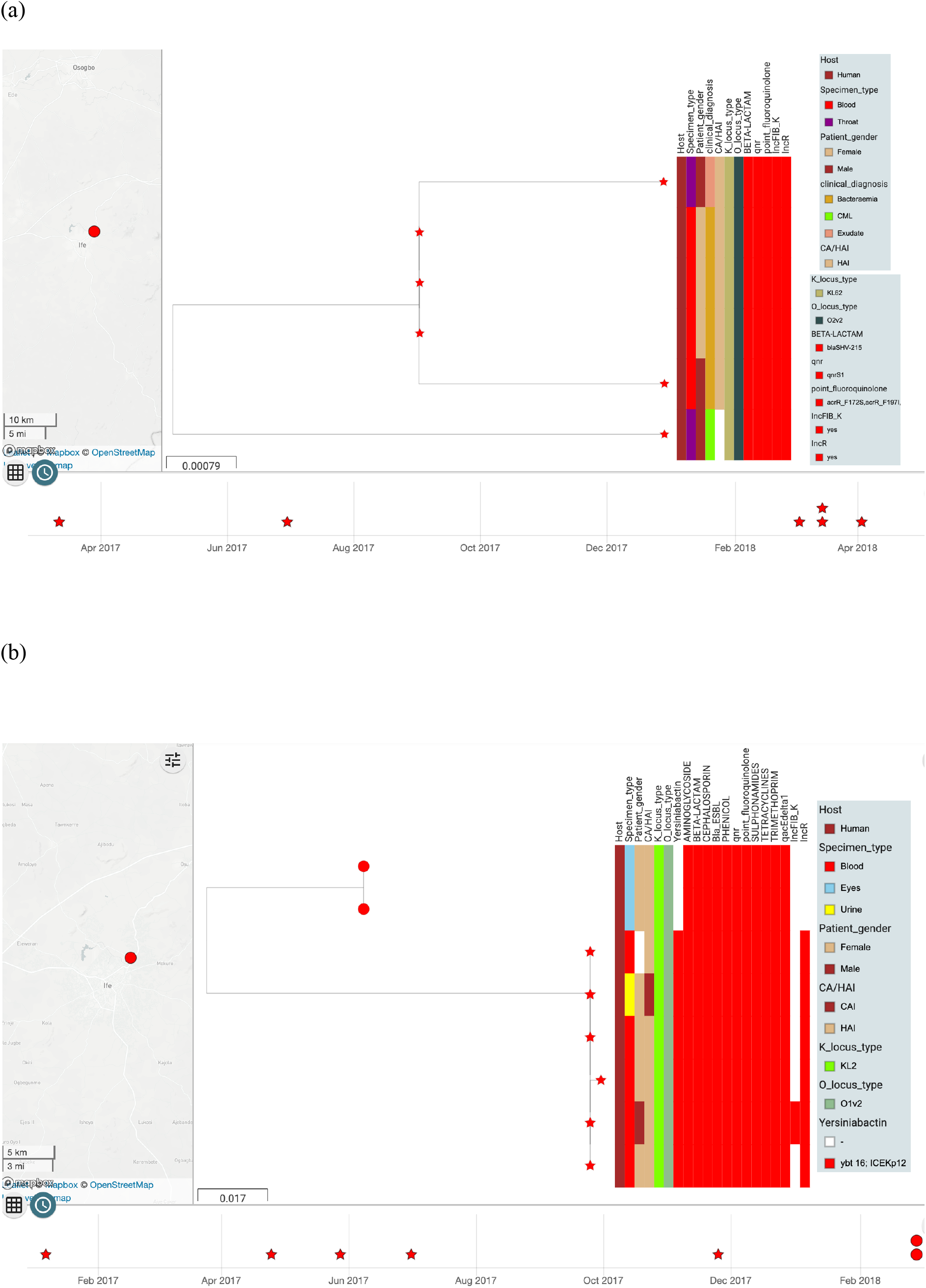
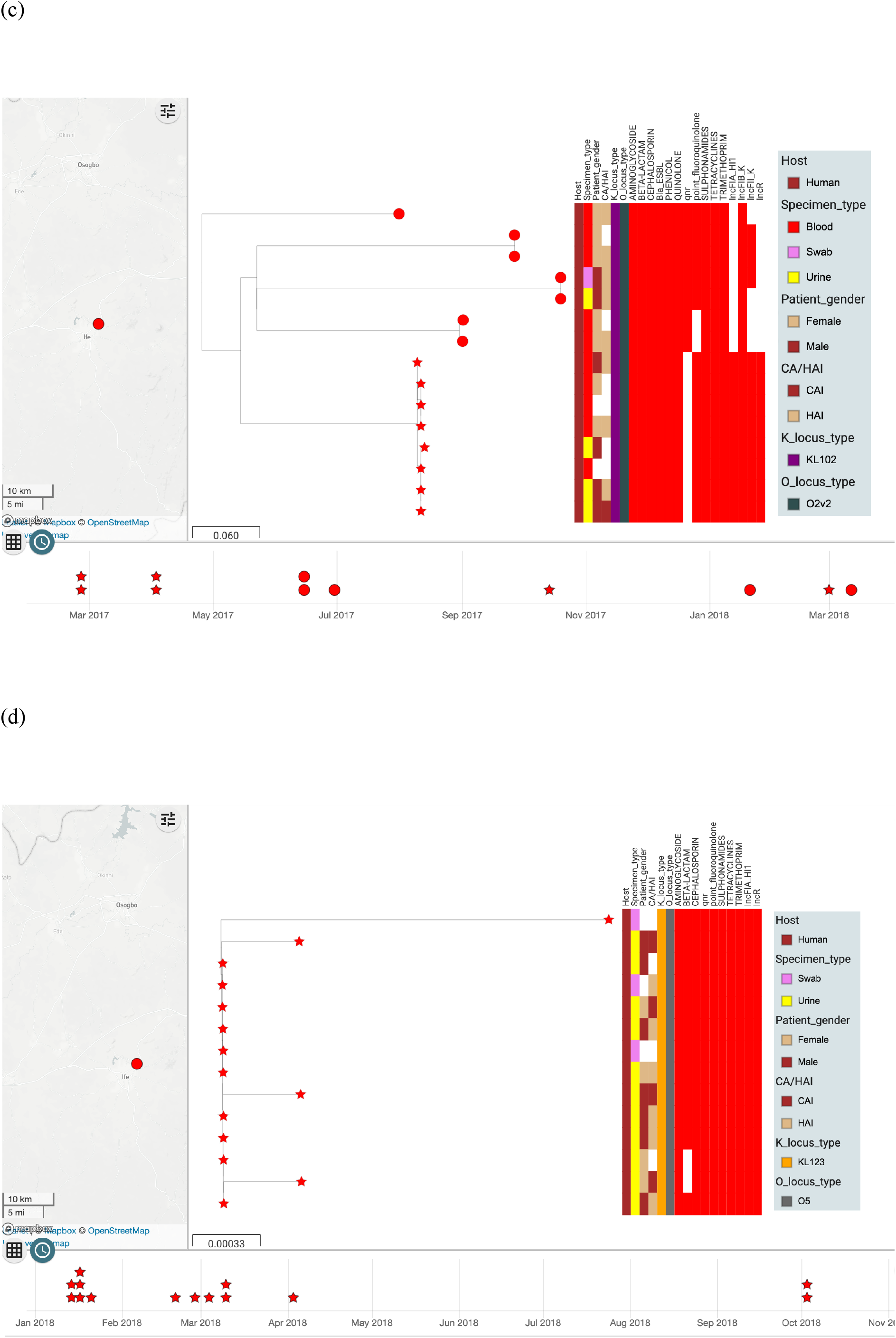
The timeline of likely outbreak of *K. pneumoniae* clones belonging to STs 17 (a), 25 (b), 307(c), and 5241(d) from the OAU sentinel site. The shape of the tree nodes depicts the outbreak (star) and non-outbreak (circle) isolate, while the color of the tree nodes depicts the sentinel site. The data are available at https://microreact.org/project/brfq17BXzwmqptfcrNRfZR/71b2f0f1 (ST17), https://microreact.org/project/8FP4F1D5fSQMv6FDxbR39b/bd381f0b (ST25), https://microreact.org/project/sV5NsJ8szcorvFAeFQgV4E/6498edf1 (ST307), and https://microreact.org/project/u5RPX7CjyitjRYMQ9ByW4W/1a3570e2 (ST5241).

## DISCUSSION

This report on the population structure of *Klebsiella* in 3 tertiary hospitals addresses critical knowledge gaps regarding the characteristics of *K. pneumoniae* in Nigeria and presents 5 *K. quasipneumoniae* genomes, which without WGS cannot be differentiated from *K. pneumoniae* in our setting, even at reference laboratory level. Ours and other data reveal that multiple *K. quasipneumoniae* lineages, including resistant ones, circulate in Nigeria [9]. We also corroborate previous reports on the diversity of *K. pneumoniae*, which pose challenges as well as opportunities for curtailing and combating this pathogen [29, 30].

The high prevalence genes conferring resistance to fluoroquinolones and extended-spectrum β-lactams, last-line options for Nigeria’s least affluent patients, in the most common lineages, reported here is of concern in spite of relatively low rates of carbapenem resistance in our setting, or elsewhere in Africa, compared to other countries [15, 31, 32, 33]. Moreover, while uncommon in this study, ST147 and ST392 strains bearing *bla_NDM-1_*, and ST101 strains carrying *bla_OXA-48_*were documented [37]. All these lineages deserve close and careful monitoring in national prospective surveillance, and further work is necessary to identify and track mobile elements they carry. The preponderance of plasmids in the *Klebsiella* strains sequenced reflects the known propensity of this genus to harbor and diversify mobile elements in the face of high selective pressure and has implications for resistance in the many species to which *K. pneumoniae* transmits DNA [2, 34].

Clonal group (CG) 307 is believed to be displacing the notorious CG258 as a primary lineage in South Africa, Italy, Colombia, and the USA, and it may be more virulent [31, 36–40]. The high ST307 frequency here, and in Malawiand South Africa, may explain the low overall prevalence of carbapenem resistance we observed compared to other locations [14, 17, 32, 35]. We recorded 21 ST307 isolates, but no ST258 isolates, and only 7 CG258 isolates ––5 ST340 and 2 ST11. Clusters of likely clonal isolates within clonal complexes 15, 25, 17, ST5241, 392, and 101 were identified among retrospective isolates from OAU, the most sampled sentinel site, within clonal complexes, 15, 25, and 17, as well as ST307, ST5241, 392, and 101. The close genetic distances observed within 4 of these clones (< 4 pairwise SNP differences in each case) and the similar or identical resistance, virulence, and plasmid replicon profiles, and temporal clustering of their isolation strongly suggest that each of these clusters represents a retrospectively identified outbreak. As fewer retrospective isolates were stored at the other two institutions, we cannot rule out similar occurrences that are below our limits of detection. Outbreaks were caused by highly resistant lineages such as the ST17 cluster, but also more sensitive ones like the ST5241 and ST307 *K. pneumoniae* clusters. While outbreaks can sometimes be detected by phenotypic testing in diagnostic labs alone, the minimal biochemical and narrow disc diffusion test repertoires employed at sentinel labs, which underlie many species’ misidentifications (Supplementary Table 1), make detection unlikely when the number of outbreak isolates is small. Our data show that, as in countries with more intensive surveillance, outbreaks likely commonly occur within health facilities in Nigeria and elsewhere in West Africa and will typically be missed without genomic support [7, 12, 42, 43]. OAU and UCH currently provide subsidized blood culture for many patients, but most Nigerian facilities with blood culture do not, and health insurance coverage is low. Therefore, identifying outbreaks is likely to be curtailed by patients’ inability to pay. Our data reveal that recent efforts to strengthen Nigeria’s AMR response and boost infection prevention and control could be mutually enhancing if blood culture of routine febrile patient blood culture is facility-supported, and if prospective genomic surveillance is used.

This study has some limitations. First, very few archival isolates from only 3 facilities were available, and epidemiological data were incomplete. Furthermore, the sequence types observed here were sampled only from invasive infections. Nonetheless, this study represents an important starting point for documenting *Klebsiella* lineages, and we will build on these findings with prospective surveillance at more hospitals in southwestern Nigeria, which have now been enrolled in the national surveillance system.

## CONCLUSION

We have detailed the characteristics of *Klebsiella* isolates from 3 southwestern hospitals in Nigeria. In addition to incorporating Nigeria-derived data into global phylogeographies of multidrug-resistant *K. pneumoniae*, our findings point to the significance of ST307 in Nigeria. Carbapenem-resistant STs 147 and 392, as well as novel ST5241 *K. pneumoniae*, may also represent future surveillance priorities for southwestern Nigeria. We have shown the benefits of public health genomic surveillance of pathogens –– in particular, the potential for outbreak identification. Infection prevention and control is a key pillar of Nigeria’s AMR National Action Plan and the data from this paper suggest that genomic surveillance and IPC implementation could be mutually reinforcing in this collaboration among scientists, health institutions, and the public health authority in Nigeria.

## ABBREVIATIONS

OAU: Obafemi Awolowo University Teaching Hospital
NCDC: Nigeria Centre for Disease Control
LAU: Ladoke Akintola University Teaching Hospital
UCH: University College Hospital
GLASS: Global Antimicrobial Resistance Surveillance System
AMR: Antimicrobial resistance
ENA: European Nucleotide Archive
WGS: Whole Genome Sequencing
CLSI: Clinical & Laboratory Standards Institute
ESBL: Extended Spectrum beta-lactamase

## ACKNOWLEDGMENTS

Members of the NIHR Global Health Research Unit for the Genomic Surveillance of Antimicrobial Resistance: Khalil Abudahab, Harry Harste, Dawn Muddyman, Ben Taylor, and Nicole Wheeler of the Centre for Genomic Pathogen Surveillance, Big Data Institute, University of Oxford, Old Road Campus, Oxford, United Kingdom and Wellcome Genome Campus, Hinxton, UK; Pilar Donado-Godoy, Johan Fabian Bernal, Alejandra Arevalo, Maria Fernanda Valencia, and Erik C. D. Osma Castro of the Colombian Integrated Program for Antimicrobial Resistance Surveillance – Coipars, CI Tibaitatá, Corporación Colombiana de Investigación Agropecuaria (AGROSAVIA), Tibaitatá – Mosquera, Cundinamarca, Colombia; K. L. Ravikumar, Geetha Nagaraj, Varun Shamanna, Vandana Govindan, Akshata Prabhu, D. Sravani, M. R. Shincy, Steffimole Rose, and Ravishankar K.N of the Central Research Laboratory, Kempegowda Institute of Medical Sciences, Bengaluru, India; Jolaade J Ajiboye of the Department of Pharmaceutical Microbiology, Faculty of Pharmacy, University of Ibadan, Oyo State, Nigeria; Celia Carlos, Marietta L. Lagrada, Polle Krystle V. Macaranas, Agnettah M. Olorosa, June M. Gayeta, and Elmer M. Herrera of the Antimicrobial Resistance Surveillance Reference Laboratory, Research Institute for Tropical Medicine, Muntinlupa, the Philippines; Ali Molloy, alimolloy.com; John Stelling, The Brigham and Women’s Hospital; and Carolin Vegvari, Imperial College London.

We sincerely appreciate technical assistance from sentinel site staff as well as Ifeoluwa Akintayo, Ifeoluwa Komolafe, and Faith Ifeoluwa Popoola. We also thank the team of curators of the Institut Pasteur MLST/cgMLST databases for curating the data and making them publicly available at https://bigsdb.pasteur.fr. We thank Lucy Goodchild for editorial assistance.

